# Maximizing citizen scientists’ contribution to automated species recognition

**DOI:** 10.1101/2022.02.17.480847

**Authors:** Wouter Koch, Laurens Hogeweg, Erlend B. Nilsen, Anders G. Finstad

**Affiliations:** Department of Natural History, Norwegian University of Science and Technology, Trondheim, Norway; Norwegian Biodiversity Information Centre, Trondheim, Norway; Intel Benelux, High Tech Campus 83, 5656 AE Eindhoven, The Netherlands; Naturalis Biodiversity Center, PO Box 9517, 2300 RA, Leiden, The Netherlands; Faculty of Biosciences and Aquaculture, Nord University, Steinkjer, Norway

**Keywords:** Image recognition, Taxonomic bias, Value of Information, Citizen science

## Abstract

Technological advances and data availability have enabled artificial intelligence-driven tools that can increasingly successfully assist in identifying species from images. Especially within citizen science, an emerging source of information filling the knowledge gaps needed to solve the biodiversity crisis, such tools can allow participants to recognize and report more poorly known species. This can be an important tool in addressing the substantial taxonomic bias in biodiversity data, where broadly recognized, charismatic species are highly overrepresented. Meanwhile, the recognition models are trained using the same biased data, so it is important to consider what additional images are needed to improve recognition models. In this study, we investigated how the amount of training data influenced the performance of species recognition models for various taxa. We utilized a large Citizen Science dataset collected in Norway, where images are added independently from identification. We demonstrate that while adding images of currently under-represented taxa will generally improve recognition models more, there are important deviations from this general pattern. Thus, a more focused prioritization of data collection beyond the basic paradigm that “more is better” is likely to significantly improve species recognition models and advance the representativeness of biodiversity data.

## Introduction

Addressing the current crisis related to the loss of biodiversity necessarily involves addressing several fundamental knowledge gaps^1,2^. Currently there are vast spatial, temporal and especially taxonomic gaps and biases in global primary biodiversity data sets, and these biases are limiting our understanding of the earth’s biosphere^3–6^. Automatic or semi-automatic observation methods based on image recognition hold great promise in solving some of the taxonomic biases currently experienced^7^. This can involve various observation methods ranging from remotely operated vessels to camera traps and citizen science programs^8–10^. Citizen science (observations made by non-professional volunteers^11^) has emerged as a very large source of biodiversity data with the potential to fill gaps in our current knowledge about the occurrence of species in time and space^12–14^. Several citizen science programs, e.g. iNaturalist, eBird, iSpot, etc.^15^ contribute data on vast scales and in amounts that cannot feasibly be acquired in any other way, with the added benefit of educating and engaging the general public^16–18^. Some of the main concerns related to citizen science data are reliability of the taxon identifications reported^19,20^, and the over-representation of more charismatic species^21–23^. Improving the quality of citizen science data is thus a vital step in addressing the knowledge gaps in our understanding of the earth’s biosphere.

Machine learning tools can help citizen scientists recognize more species and provide a quality control mechanism that helps to reduce the risk of species misidentification^7^, but their performance is inherently linked to the quality of the data used to train them. Such tools are increasingly used to help citizen scientists identify species from images^24–26^, and in doing so, help address the aforementioned issues in citizen science data to some degree by providing an independent way to verify identifications, and helping citizen scientists report a broader range of taxa than they would otherwise be familiar with^27^. Observations accompanied by images can be used for training an image recognition model for use in the field, and generally one would use images provided by citizen scientists as training data for models to be used by citizen scientists, as the context and general type of such images is most similar to the intended use. Deep neural networks are designed to draw inferences from novel data by generalized patterns observed in training data^28^, and as such generalization requires a lot of training data, they in turn depend heavily on the data corpus citizen science contributions provide. In this manner, citizen science and automated image recognition are increasingly interdependent, with image recognition models helping citizen scientists collect data to expand our knowledge base, whilst themselves also depending on the collection of more images in order to improve the next generation of recognition models.

While some species are readily recognized with limited experience, others require extensive experience with many specimens to obtain the necessary knowledge. Machine learning is no different from human learning in this respect; different amounts of training data are required depending on the distinctness of species’ characteristics. Therefore, there can be substantial differences between taxa in the number of images required per species for the best achievable model performance, depending on species’ distinctiveness, the variation in appearance, the various angles and contexts in which photos are taken, the extent to which a species’ behavior is suited for high quality documentation, etc.^27,29,30^ As a result, the marginal value of adding a new image to the training set is not equal across taxa, but varies both because the size of the existing training set is different, and the fact that some species are more distinct than others. Thus it is important to consider the informational value of added citizen science observations with images.

In this study, we use a large Norwegian citizen science project as an example to investigate the nature of the bias in citizen science image data, and how this relates to the value of data for image recognition models. One way to evaluate this is by using the concept of Value of Information; “*the increase in expected value that arises from making the best choice with the benefit of a piece of information compared to the best choice without the benefit of that same information*”^31^. Considering training data for image recognition models for citizen science in the Value of Information framework allows us to identify the most effective prioritization of data collection for improving recognition models, rather than simply adding more for all taxa, or more for taxa that are currently the most under-represented. First, we evaluate whether the biases generally found in observation data are the same within citizen science observations with images, or if there are different biases that need to be taken into account. Then we train multiple image recognition models for different taxa, with a gradually increasing number of images per species, allowing us to quantify and compare the effects of adding more training data between taxa. Using these performances, we estimate the Value of Information of adding training data for each taxon, relative to the amount of images that are currently available. Finally, comparing this Value of Information to the amount of over- or under-representation of these taxa, we demonstrate that mobilizing images with a higher Value of Information provides an alternative, data-driven and efficient approach compared to simply prioritizing images of the currently most under-represented taxa.

## Results

### Taxonomic bias in Citizen Science observations with or without images

It has been well documented in a global context that particularly charismatic taxonomic classes have many times more reported observations per species than those that are considered less charismatic^5^. We find the same pattern when considering classes within the totality of GBIF mediated observations for Norway from all sources (figure 1a). When limiting this analysis to only observations with images that originate from the citizen science platform Species Observation Service^32^, a different pattern emerges (figure 1b). Perhaps most eye-catchingly, Insecta are the most under-represented taxon in the totality of Norwegian observations, but the 3rd most over-represented when limiting the analysis to citizen science images. We performed a similar analysis for the 12 taxonomic orders used in the machine learning part of this study (figure 4), revealing biases in relative presentation per species at this finer taxonomic scale for citizen science observations with images; the data available for training out recognition models.

**Figure 1:**
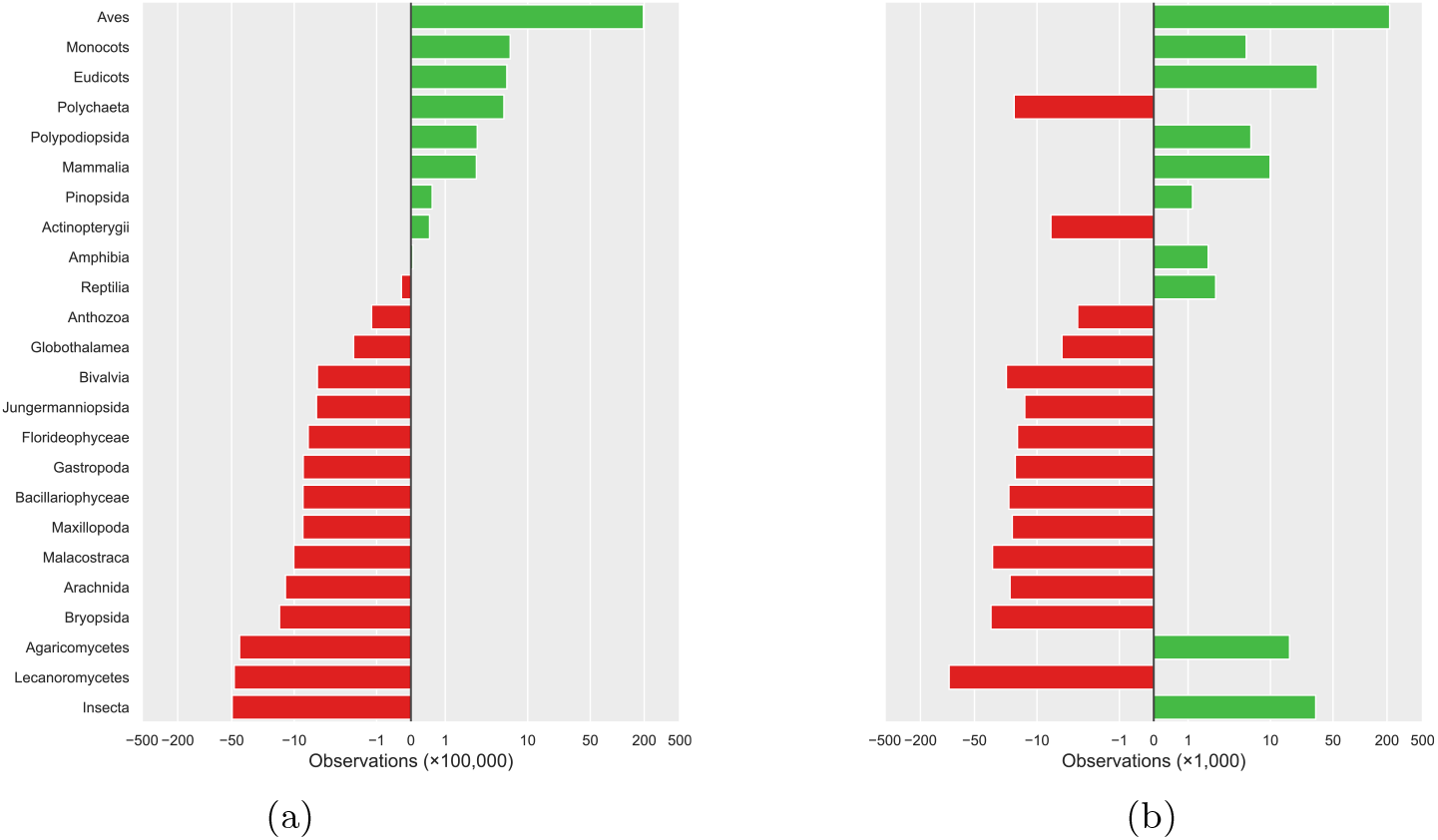
The per-species representation of observations in Norway per class, using all GBIF data (a) or only GBIF mediated citizen science data with images (b). The 0-line is where the values would be if the average number of observations per species in that class was equal to the average number of observations per species over all classes combined. Plotted here on an inverse hyperbolic sine-transformed scale, sorted by the per-species representation in subplot (a).

### Image recognition performance and the Value of Information

When training image recognition models, the amount of training data provided to the model determines how well the model is able to recognize species in the test images. For all orders, as models are provided with more images per species, their performance (as measured by the F_1_ scores) increases. Comparing the performances for each order with the lowest and highest number of training images per species, as well as the gradual performance increase over intermediate numbers of training images, it is clear that the 12 orders have distinct performance curves (figure 2). From this it follows that the increase in performance at any given point on these curves - the Value of Information of adding images at that point - also differs between orders. Combined with widely different amounts of currently available observations between orders, the estimated VoI of an image added to the images that are currently available for that order also varies widely (figure 3).

**Figure 2:**
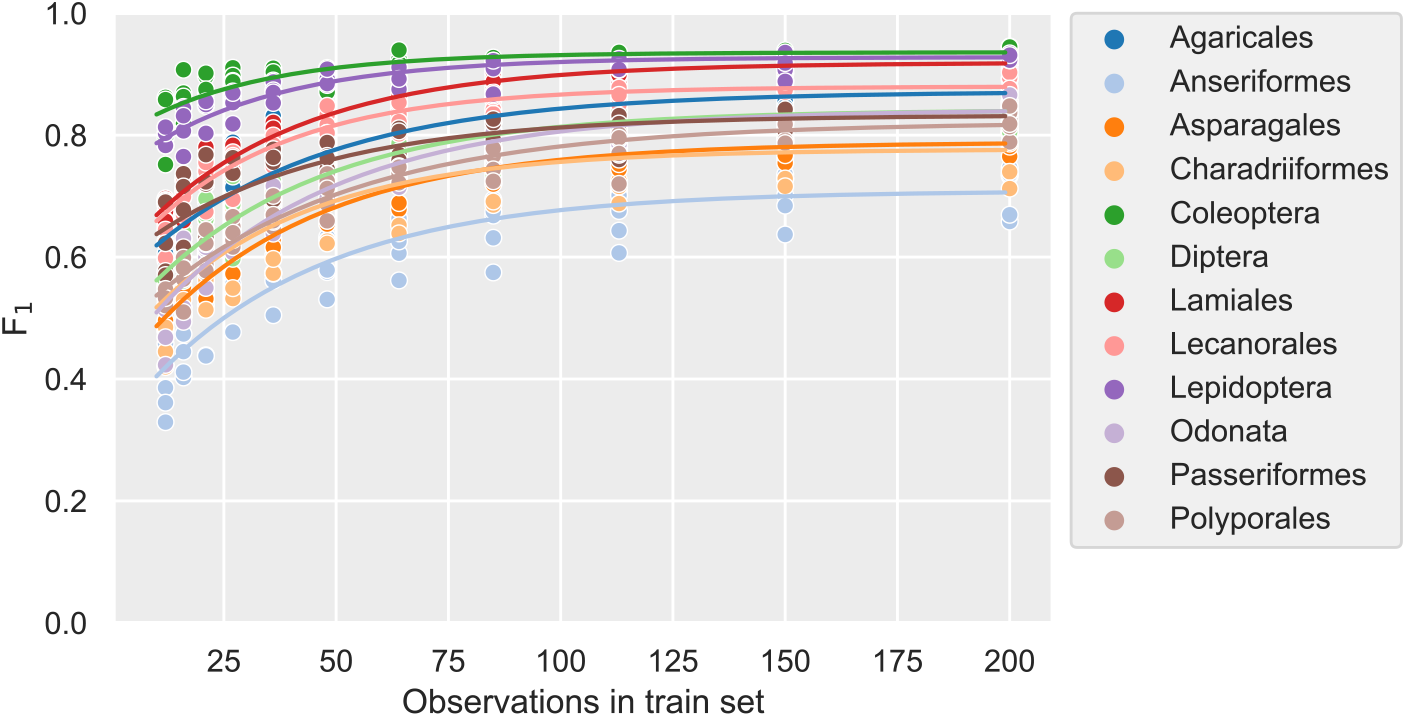
The performance (F_1_ score) vs the train set size. Lines are the fitted Von Bertalanffy Growth Function-curves per order. See the Supplementary Information for an interpretation of such curves.

**Figure 3:**
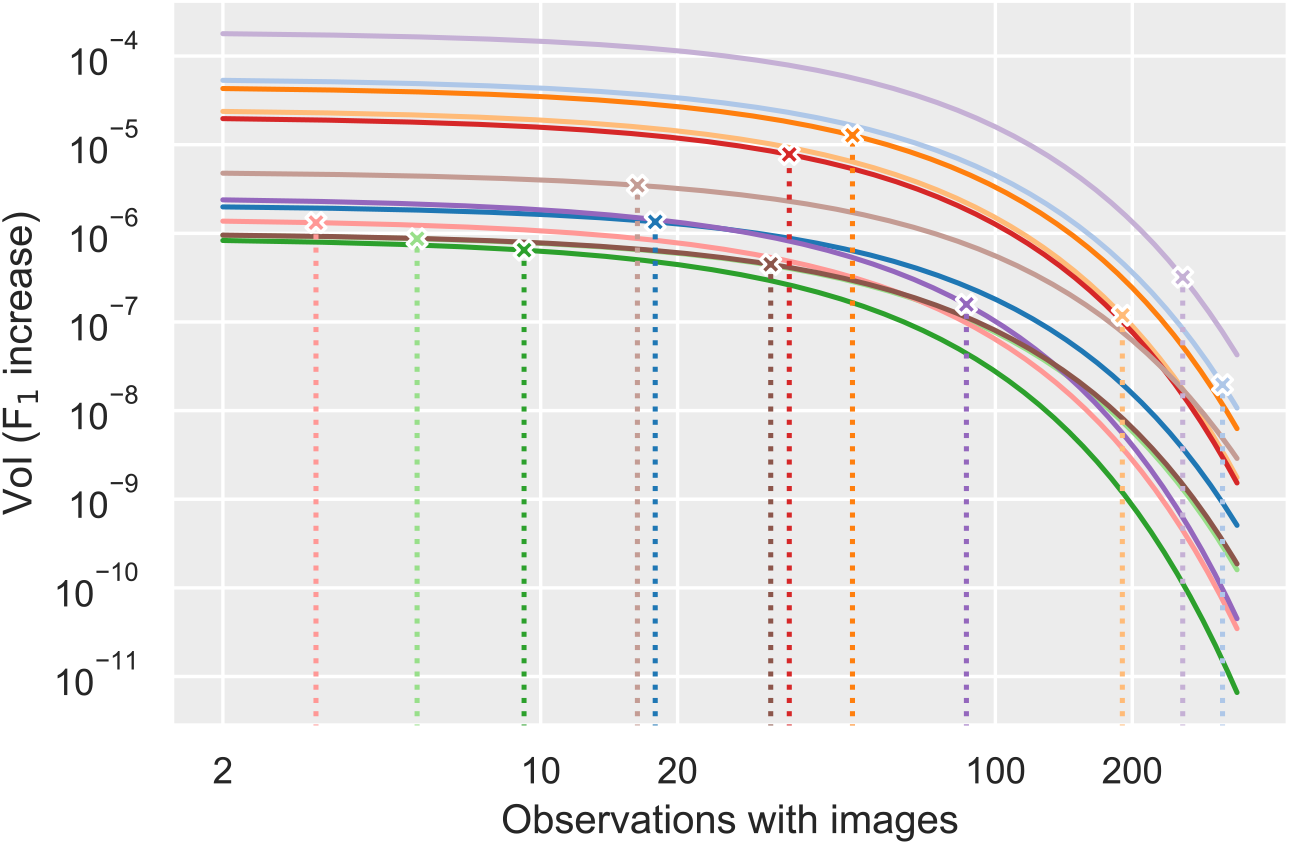
The VoI (F_1_ increase) for each order as the result of adding a single image for a single species, versus the average number of images available per species. Dotted lines mark the average number of images per species currently available for the respective order, from which the current expected VoI (marked with x) is derived. Colors denote each order as in figure 2.

### Combining VoI and taxonomic bias

After obtaining the per-species over- and under-representations, as well as the current expected Value of Information of additional images, we can compare the two values for each order in the experiment. Plotting the taxonomic bias of the orders used in this experiment together with their estimates for their respective estimated VoI, it is clear that current under- or over-representation of the order is not the determining factor for the expected value of additional observations. While the VoI of under-represented orders is generally higher, differences between orders in their learning curves cause some orders to have a higher or lower VoI than just their overall over- or under-representation would indicate (figure 4).

**Figure 4:**
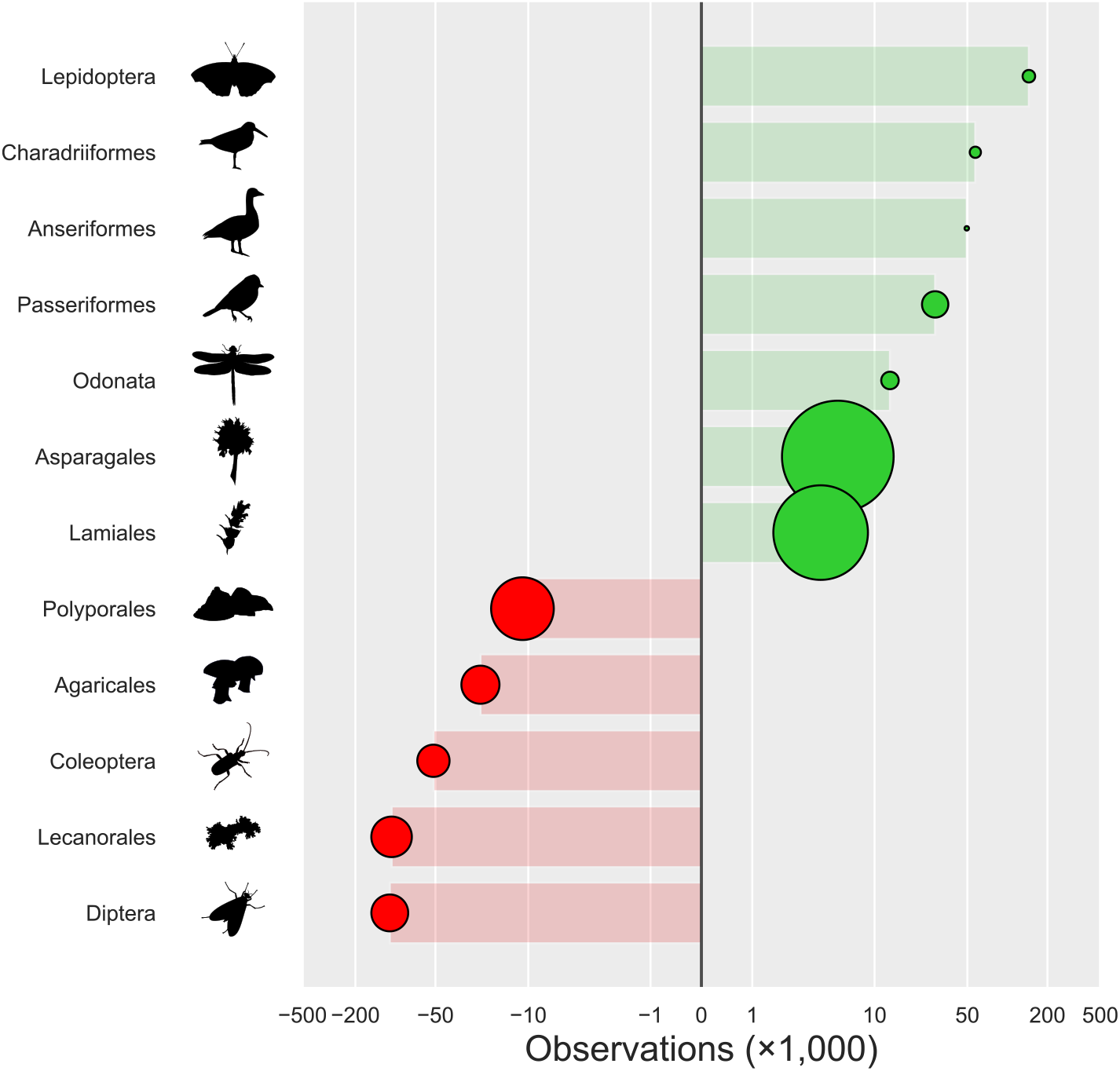
The relative per-species representation in Norwegian citizen science observations with images, and their Value of Information (VoI). The areas of the circles are relative to their respective VoI, defined as the current expected performance increase (in F_1_ score) for one added observation with images for that order. If the VoI of adding data was mainly determined by the current relative over- or under-representation of a taxon, one would expect circles to gradually increase for more under-represented orders in the lower part of the graph. Numerical values provided in the Supplementary Information.

## Discussion

We set out to investigate the taxonomic bias in citizen science data, in particular when accompanied by images, using a large Norwegian citizen science project as an example case. Such images can be used to train deep neural networks for image recognition, helping citizen scientists by verifying species identifications and address some of the inherent taxonomic bias. By examining how the performance of recognition models increases as it is provided with more images in an experimental setup, we can estimate how much we expect models to improve when adding more images to those currently available for each taxon. Comparing this Value of Information to the taxonomic bias within citizen science image data, we propose data prioritization strategies based on what additional data would improve recognition models the most. Such strategies would be more efficient than merely focusing more on taxa for which there are currently fewer images available.

### Taxonomic bias

The taxonomic biases within citizen science observations considered in the current study follow a similar pattern to what has been found across biodiversity data in general^5^. However, when only considering citizen science observations with images, these trends are less pronounced; plants and fungi have relatively higher percentages of observations with images than for example birds (figure 1b). This indicates that while birds are still the most reported group also within citizen science observations with images, bird observations are generally less commonly documented with images. The reverse is true for the Insecta, which are so abundant in the citizen science image data as to be the 3rd most overrepresented class in that context. This is in stark contrast to what has been found for the totality of GBIF mediated observations globally^5^ and in the Norwegian context we examined here, where the Insecta are the single most under-represented class.

Analyzing the taxonomic biases for the orders used in the machine learning part of this study sheds some light on the underlying mechanisms. While all orders within Aves are over-represented regardless of the nature of the observations considered, the Insecta are more diverse in their bias.

We hypothesize that this disparity between taxonomic bias in all data versus that in citizen science data with images is most likely a combination of the behavior of the species and the kind of citizen scientists reporting the observations. There are distinctly different types of citizen scientists, with their own contribution patterns^33^. For casual reporters lacking specialized equipment, charismatic butterflies and flowering plants are more readily photographed opportunistically than birds. Meanwhile, a group of quite persistent ornithologists report the bulk of the bird observations in the dataset. This is typically a group reporting in a structured manner, more often based on local inventories and checklists, where reporting with images is less common than with opportunistic observations.

### Image recognition and Value of Information

There are clearly differences between orders in the rates at which image recognition improves as more images are made available per species (figure 2). These differences between orders manifest in both initial performances, the rate at which performances change, and the maximum performance achieved. This indicates that, as is the case for humans, it requires more experience to learn to identify species within certain taxa than others, while the reliability with which species are correctly identified once the necessary knowledge has been acquired also differs. The differences between orders in this regard is not necessarily directly linked to the taxon’s characteristics alone, however. Image quality and composition can vary between taxa depending on factors such as specimens’ behavior or lack thereof, physical size, and the kind of citizen scientist generally photographing the species.

The VoI estimates for each of the orders provides equally diverse results. For any given number of images per species, orders differ in the expected performance increase at that point, as do the relative rates at which these performances change as data is added. As a consequence, there is a range of varying estimates for the VoI for each order, depending on both the number of images currently available per species, and the way the VoI per additional image declines as more images are already available to the model.

### Combining taxonomic biases and the Value of Information

As we have an estimate of how over- or under-represented the orders with which the recognition models have been trained are relative to one another, as well as a per-order estimate of the VoI per added image, we can address the question whether models are best improved by adding more image data equally across orders, if one should ideally prioritize under-represented orders, or if there is a prioritization to be made based on order-specific differences. As shown in figure 4, there are distinct differences in the Values of Information per order, and these do not merely correlate with their respective over- or under-representation. The plant orders of Asparagales and Lamiales clearly have a higher VoI despite their slight over-representation when compared to the other orders in this experiment. The fungi order Polyporales also gains more than twice the VoI per additional image in comparison to the fourth-most valuable order, the Lecanorales. We conclude that, from a VoI perspective, these are the orders for which a recognition model would benefit the most per image added, also when compared to other, more under-represented orders.

## Conclusions

Based on the Value of Information for image recognition models, a citizen scientist or citizen science project manager wishing to maximize their impact in this regard might want to focus on orders with the highest expected VoI per image added, rather than simply on the order with the lowest number of images per species. Observations with images of other orders, while in some cases less well-represented in the available image data, appear to provide less VoI per observation added. As citizen scientists are in large part motivated by a desire to advance scientific knowledge^34^, communicating such considerations can be an important part of community engagement.

In generalizing these findings, the following has to be noted:

- The taxa identified here as having the highest expected VoI per image added are examples from the limited subset of orders used within this experiment. As illustrated by the observed variation in per-species representation and VoI between orders that belong to the same class, it is evident that generalization of a class like Insecta fails to give insight into intra-class variation. It is likely that a similar principle applies to orders, where for example a taxonomic group like Norwegian warblers likely has a different VoI curve than the more readily distinguished titmice. Such differences will remain hidden from view when analyzing passerine birds as a single taxonomic group.
- Our findings are derived from Norwegian species reported on a single Norwegian citizen science portal. The diversity of species within the same orders can differ in other regions, affecting the VoI curves. Different portals will also differ in the way they accommodate reporting observations with images, and in general attract different types of users^23^. All of this is likely to have an effect on the proportion of observations accompanied by photographic evidence and the quality thereof. Such factors also affect the nature of newly added data, including its expected VoI.
- Models were trained on species for which at least 220 observations with images were available. This is not a random subset of all the species within an order, and likely to be biased towards charismatic species and those that are more readily identifiable from an image. This can lead to an overestimation in terms of learning rate and thus the VoI curve, especially within orders in which relatively few species have the data availability we selected for here. Then again, future observations to be added to the data will be prone to the same biases, in which case the VoI of such an addition will be lower than it would be for a truly random species.

Regardless of the specific taxa and derived values, our findings demonstrate that a more informed decision is possible when choosing to focus on certain taxa for data collection. Prioritization of taxa for which to mobilize additional data can be informed by considering its expected Value of Information, rather than simply prioritizing those that are currently the most under-represented numerically.

Training machine learning models requires a lot of data, certainly when context, morphology and phenology vary, such as when classifying in situ images. Data collection in machine learning generally is a matter of harvesting whatever one can to provide the model with more data. Within (citizen) science, the collection of images mainly serves as secondary data, providing documentation for the occurrence it accompanies. With the more widespread use of image recognition models as both a user tool and a mechanism for quality control, it is time to view images as data in and of themselves. Such a shift calls not only for conscious choices when it comes to the Value of Information in images, but increased implementation of data practices such as persistent storage, metadata standardization and the other FAIR data principles^35^ to enable more apt usage of image data for current and novel applications.

## Methods

In the current study we utilize an extensive network and data from Citizen Science in order to test for among taxa variation in biases and Value of Information in image recognition training data. We use data from Norway as an example dataset due to the complete spatial and taxonomic coverage of the citizen science data, reported both with and without images. An advantage of this particular citizen science project is that no image recognition model has been available to the reporters and that data have been bulk-verified by experts. This ensures that the models trained in this experiment are not trained on the output resulting from the use of any model, but with identifications and taxonomic biases springing from the knowledge and interest of human observers. Moreover, the authoritative Norwegian taxonomy allows for analyses on taxonomic coverage.

In an exploration procedure we determined the taxonomic level of orders to be suitable examples of taxa with a sufficiently wide taxonomic diversity, and enough data in the dataset to be evaluated for models in this experiment. For the selected orders, as well as the classes used by Troudet *et al*.^5^, we acquired taxon statistics and observation data from the Global Biodiversity Information Facility, GBIF, the largest aggregator of biodiversity observations in the world^36^. The authoritative taxonomy for Norway was downloaded from the Norwegian Biodiversity Information Centre. In the experimental procedure, models were trained for 12 distinct orders, artificially restricting these models to different amounts of data. In the data analysis stage, model performances relative to the amount of training data were fitted for each order, allowing the estimation of a Value of Information. Using the number of observations per species on GBIF, and the number of species known to be present in Norway from the Norwegian Species Nomenclature Database, we calculated relative taxonomic biases.

### Exploration

Initial pilot runs were done on 8 species groups, using different subset sizes of observations for each species, and training using both an Inception-ResNet-v2^37^ as well as an EfficientNetB3^38^ architecture for each of these subsets. These initial results indicated that the Inception-ResNet-v2 performance was more robust and generally higher, so subsequent experiments were done using this architecture. The number of observations which still improved the accuracy of the model was found to be between 150 and 200 in the most extreme cases, so the availability of at least 220 observations with images per species was chosen as an inclusion criteria for the further experiment. This enabled us to set aside at least 20 observations per species as a test dataset for independent model analysis.

From a Darwin Core Archive file of Norwegian citizen science observations from the Species Observation Service with at least one image^32^, a per species tally was generated. We then calculated how many species, with a minimum of 220 such observations, would, at a minimum, be available per taxon if a grouping was made based on each taxon rank level with the constraint of resulting in at least 12 distinct taxa. For each taxonomic level, we calculated how many species having at least 220 such observations were available per taxon when dividing species based on that taxon level. When deciding on the appropriate taxon level to use, we limited the options to taxon levels resulting in at least 12 different taxa.

A division by order was found to provide the highest minimum number of species (17) per order within these constraints, providing 12 orders. The next best alternative was the family level, which would contain 15 species per family, covering 12 of the 267 eligible families.

### Data collection

We retrieved the number of species represented in the Norwegian data through the GBIF API, for both all observations, all citizen science observations, and all citizen science observations with images for the 12 selected orders and the classes used by Troudet *et al*.^5^. We also downloaded the Norwegian Species Nomenclature Database for all kingdoms containing taxa included in these datasets.

Observations with images were collected from the Darwin Core Archive file used in the exploration phase, filtering on the selected orders. For these orders, all images were downloaded and stored locally.

### Experimental procedure

For each selected order, a list of all species with at least 220 observations with images was generated from the Darwin Core Archive file^32^. Then, runs were generated according to the following protocol (figure 5):

**Figure 5:**
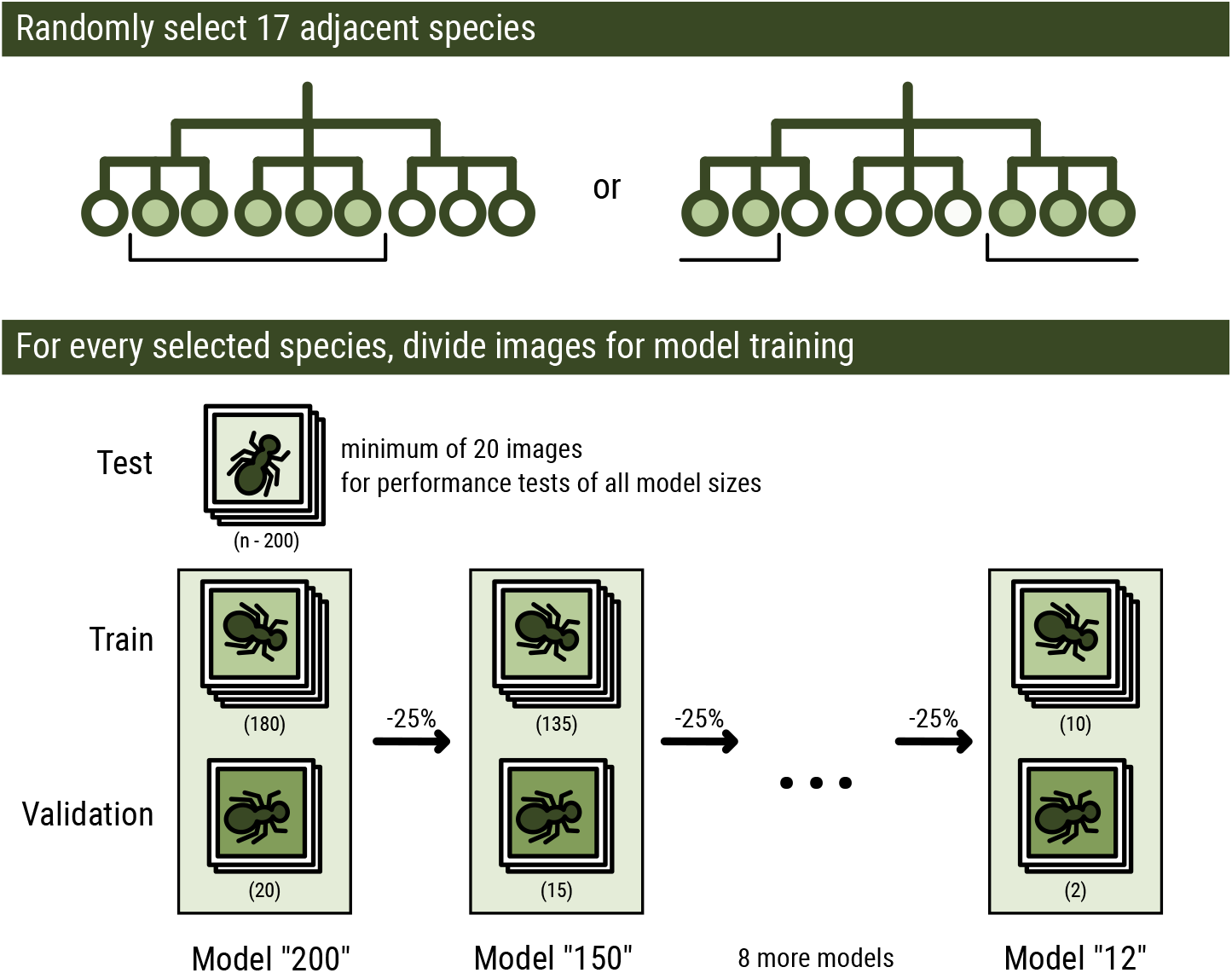
Data selection and subdivision. Each run is generated by selecting 17 taxonomically adjacent species per order, and randomly assigning all available images of each selected species to that run’s test-, train- or validation set. Training data are used as input during training, using the validation data to evaluate performance after each training round in order to adjust training parameters during training. The test set is used to measure model performance independently after the model is finalized^28^. For each subsequent model in that run, train and validation data are reduced by 25%. The test set is not reduced, and used for all models within a run.

1. From a taxonomically, alphabetically sorted list, a subset of 17 consecutive species starting from a random index was selected. If the end of the list was reached with fewer than 17 species selected, selection continued from the start of the list. The taxonomic sorting ensures that closely related species (belonging to the same family or genus), bearing more similarity, are more likely to be part of the same experimental set. This ensures that the classification task is not simplified for taxa with many eligible species.
2. Each of the 220+ observations for each species were tagged as being either test, training or validation data. A random subset of all but 200 were assigned to the test set. The remaining 200 observations were, in a 9:1 ratio, randomly designated as train or validation data, respectively. In all cases, images from the same observation were assigned to the same subset, to keep the information in each subset independent from the others. The resulting lists of images are stored as the test set and 200-observation task.
3. The 200 observations in the train and validation sets were then repeatedly reduced by discarding a random subset of 25% of both, maintaining a validation data proportion of *≤* 10%. The resulting set was saved as the next task, and this step was repeated as long as the resulting task contained a minimum of 10 observations per species. The test set remained unaltered throughout.

Following this protocol results in a single run of related training tasks with 200, 150, 113, 85, 64, 48, 36, 27, 21, 16 and 12 observations for training and validation per species. The seeds for the randomization for both the selection of the species and for the subsetting of training- and validation datasets were stored for reproducibility. The generation of runs was repeated 5 times per order to generate runs containing tasks with different species subsets and different observation subsetting.

Then, a Convolutional Neural Network (CNN) based on Inception-ResNet-v2^37^ (see the Supplementary Information for model configuration) was trained using each predesignated train/validation split. When the learning rate had reached its minimum and accuracy no longer improved on the validation data, training was stopped and the best performing model was saved. Following this protocol, each of the 12 orders were trained in 5 separate runs containing 11 training tasks each, thus producing a total of 660 recognition models. After training, each model was tested on all available test images for the relevant run.

### Data analysis

The relative representation of species within different taxa were generated by dividing the number of species present in the GBIF data for Norway within each taxon by the number of accepted species within that taxon present in the Norwegian Species Nomenclature Database^39^, in line with Troudet *et al*.^5^.

As a measure of model performance, we use the F_1_ score, the harmonic mean of the model’s precision and recall, given by

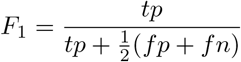

where *tp, fp* and *fn* stand for true positives, false positives and false negatives, respectively. The F_1_ score is a commonly used metric for model evaluation, as it is less susceptible to data imbalance than model accuracy^28^.

The Value of Information (VoI) can be generically defined as “*the increase in expected value that arises from making the best choice with the benefit of a piece of information compared to the best choice without the benefit of that same information*”^31^. In the current context, we define the VoI as the expected increase in model performance (F_1_ score) when adding one image-documented observation. To estimate this, for every order included in the experiment, the increase in average F_1_ score over increasing training task sizes were fitted using the Von Bertalanffy Growth Function, given by

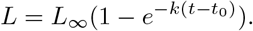

where *L* is the average F_1_ score, *L*_*∞*_ is the asymptotic maximum F_1_ score, *k* is the growth rate, *t* is the number of observations per species, and *t*_0_ is a hypothetical number of observations at which the F_1_ score is 0. The Von Bertalanffy curve was chosen as it contains a limited number of parameters which are intuitive to interpret, and fits the growth of model performance well. The estimated increase in performance at any given point is then given by the slope of this function, i.e. the result of the differentiation of the Von Bertalanffy Growth Curve, given^40^ by

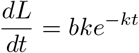

where

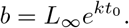

Using this derivative function, we can estimate the expected performance increase stemming from one additional observation with images for each of the species within the order. Filling in the average number of citizen science observations with images per Norwegian species in that order for t, and dividing the result by the total number of Norwegian species within the order, provides the VoI of one additional observation with images for that order, expressed as an average expected F_1_ increase.

## Supplementary Information

### Code availability

All code used in this study for the experiment and the generation of graphs provided in this manuscript is available on https://github.com/WouterKoch/citizen_science_VoI.

### Image recognition model configuration

Models were trained in Python 3.9^41^, using TensorFlow^42^ and Keras^43^ to train a new recognition model based on the Inception-ResNet-v2 architecture^37^ for every dataset. A dense classification layer using softmax activation replaced the top layer of the Inception-ResNet-v2 model as a new top layer, with 17 nodes to classify each of the 17 species. For the loss function we used standard categorical cross entropy loss.

Color channels of input images were normalized between −1 and 1, and were scaled to 256 *×* 256 pixels, cropping the image to become square if needed. Training data were augmented by shearing up to a factor of 0.2, zooming up to a factor of 0.2, rotating up to 90 degrees, and randomly flipping horizontally or not. Validation and test images were only normalized and squared, not augmented. In the first training stage, the weights of the original Inception-ResNet-v2 layers were frozen, training only the newly added top layer. This was done for 2 epochs with a learning rate of 1 · 10^*−*3^. This has an equivalent effect as learning rate warm-up.

In the second training stage, all layers were trained. This was done for a maximum of 200 epochs, with an initial learning rate of 1 · 10^*−*4^. The learning rate was multiplied by 0.1 when the validation loss did not improve for 3 consecutive epochs. The minimum of the learning rate was set to 1 · 10^*−*8^.

After each epoch, model performance was evaluated using the validation set, saving the weights of the current model to disk as the latest checkpoint if the accuracy for the validation set had improved since the last saved checkpoint. Finally, when the model did not reduce its loss for 8 consecutive epochs, training was stopped. The most recently stored checkpoint was then used as the final recognition model for that dataset, and its performance measured using the test data.

### Taxonomic order result metrics

**Table 1:**
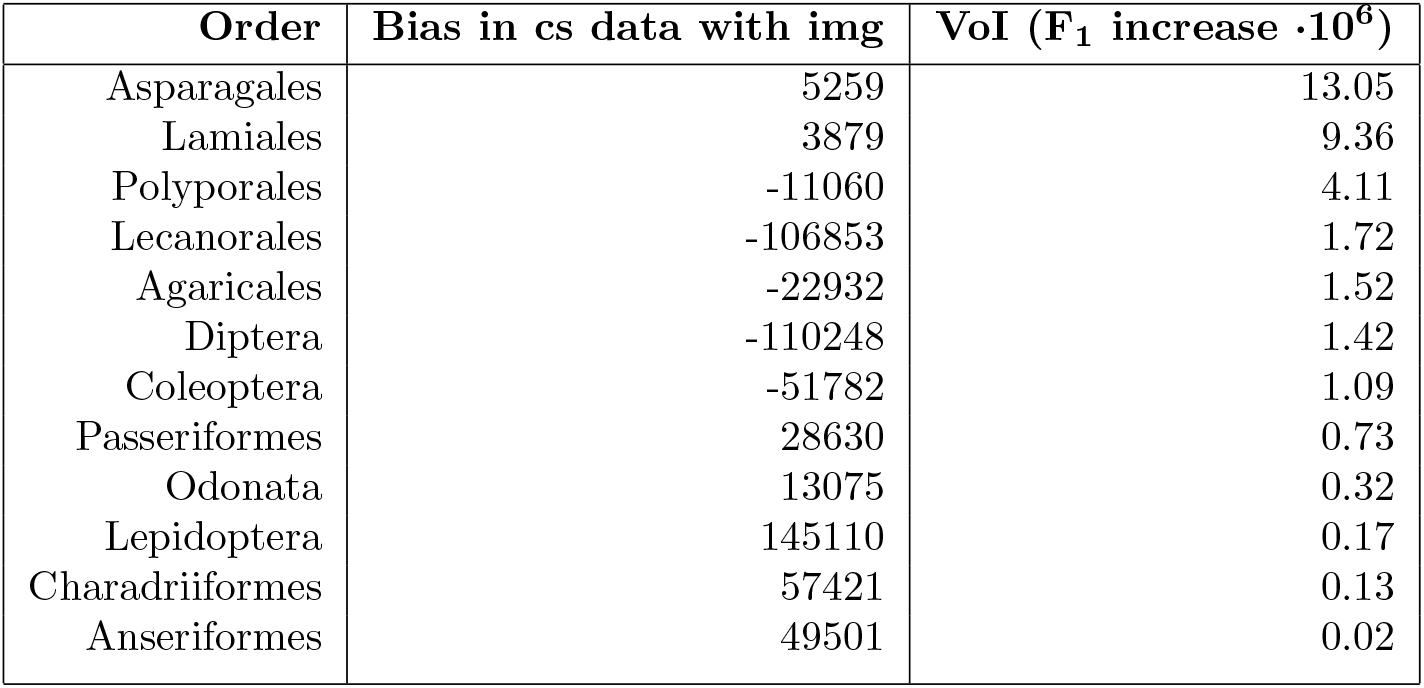
Orders used in the machine learning experiment, their over- or under-representation among citizen science observations with images (relative to all orders having an equal average amount of such observations per species), and the Value of Information as measured by the expected F_1_ increase for adding one picture to the number of images currently available. Sorted by VoI (descending). These are the numerical values for figure 4.

### Von Bertalanffy Growth Curves

**Figure 6:**
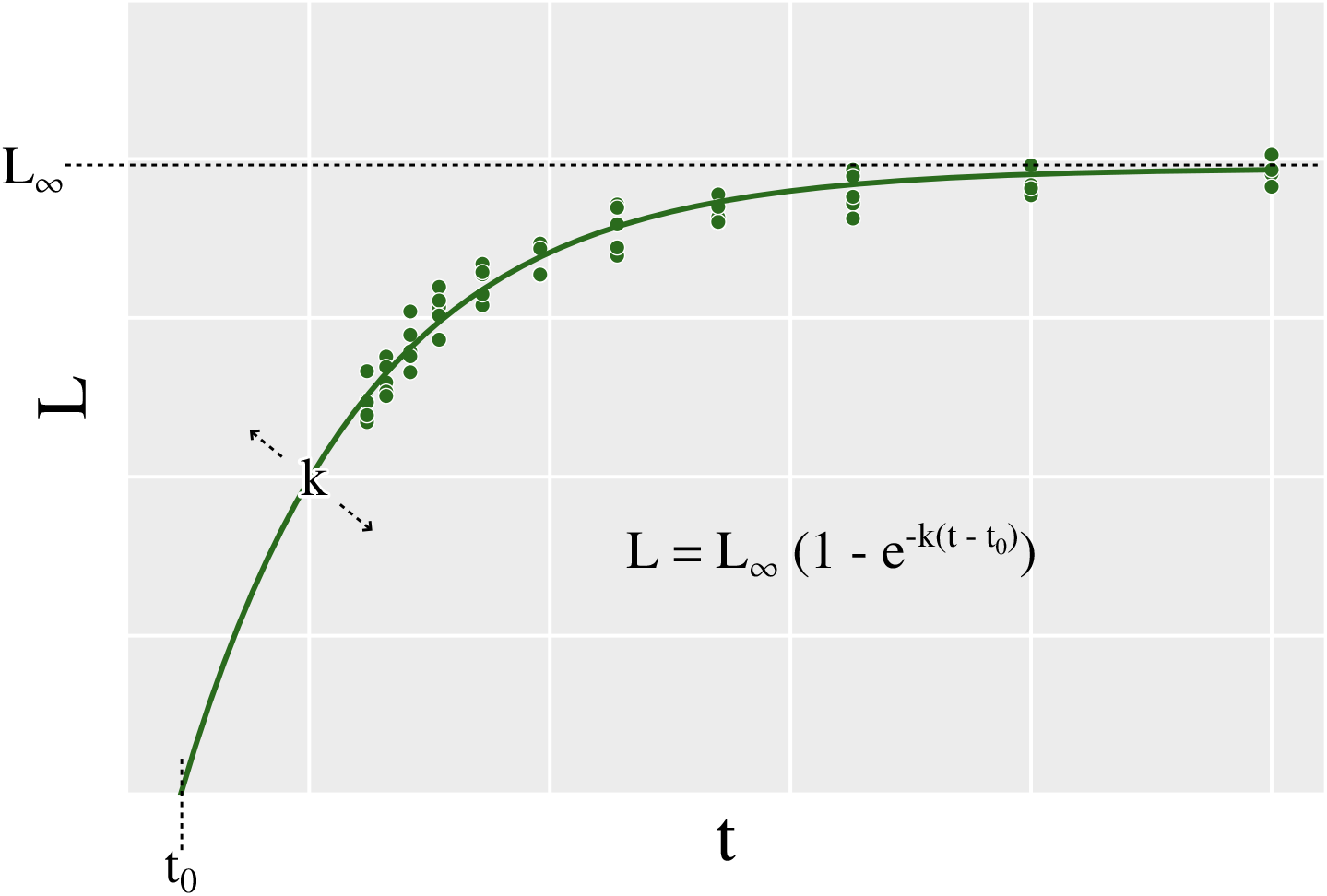
Visualization of the Von Bertalanffy Growth Curve parameters. Curves were fitted using the Levenberg-Marquardt (Least Squares) algorithm. Residuals were plotted for each taxon and not found to be heterogeneous in their distribution.

